# Insights into cerebral haemodynamics and oxygenation utilising *in vivo* mural cell imaging and mathematical modelling

**DOI:** 10.1101/195503

**Authors:** Paul W. Sweeney, Simon Walker-Samuel, Rebecca J. Shipley

## Abstract

The neurovascular mechanisms underpinning the local regulation of cerebral blood flow (CBF) and oxygen transport remain elusive. In this study we have combined novel *in vivo* imaging of cortical microvascular and mural cell architecture with mathematical modelling of blood flow and oxygen transport, to provide new insights into CBF regulation that would be inaccessible in a conventional experimental context. Our study implicates vasomotion of smooth muscle actin-covered vessels, rather than pericyte-covered capillaries, as the main mechanism for modulating tissue oxygenation. We also resolve seemingly paradoxical observations in the literature around reduced blood velocity in response to arteriolar constrictions and deduce the cause to be propagation of constrictions to upstream penetrating arterioles. We provide support for pericytes acting as signalling conduits for upstream smooth muscle activation, and erythrocyte deformation as a complementary regulatory mechanism. Finally, we caution against the use of blood velocity as a proxy measurement for flow. Our combined imaging-modelling platform complements conventional experimentation allowing cerebrovascular physiology to be probed in unprecedented detail.

## Introduction

The mammalian brain has evolved complex neurovascular coupling mechanisms to regulate the flow of blood to tissue in response to neuronal activity. However, the biophysical processes underpinning cerebral blood flow regulation are intricate and incompletely understood. For example, it is known that mural cells, which line the walls of cortical blood vessels, act as an intermediary between neural activity and blood flow, but the precise mechanisms that define this control are unclear.

*In vivo* experiments that aim to probe these mechanisms present a range of technological challenges, and typically provide limited spatial and temporal information. Increasingly, mathematical modelling is being used to provide a precisely controlled, alternative tool for investigating biophysical phenomena^8,9,12,13,23,24,34^, particularly when accompanied by corroborating *in vivo* data.

In this study, we sought to develop a novel mathematical modelling platform that used structural data from *in vivo* imaging of the mouse cortex upon which to simulate the complex physiology and fluid dynamics of the cerebral vasculature. The extraction of cerebral vascular network structure enabled the parametrisation of two comprehensive, coupled mathematical models for blood flow and oxygen transport, which facilitated physiologically realistic *in silico* experiments to be performed. These allowed us to examine the complex interactions between coupled physiological phenomena, occurring in spatially isolated regions within the cortical vasculature.

Our novel modelling approach inputs data on the location of vessels surrounded by smooth muscle cells (SMCs), which were derived from *in vivo* two-photon imaging of a genetic reporter of alpha-smooth muscle actin (*α*-SMA). SMCs are included in the four morphological classes of mural cells that are thought to have an influence on CBF regulation^19^. SMCs surround the entire circumference of pial and penetrating arterioles (with diameters in the range 15 to 40 *μ*m), precapillary arterioles (diameters ranging from 3 to 15 *μ*m) and postcapillary venules, whilst capillary pericytes are found on microvessels with diameters of 3 to 9 *μ*m. Conversely, capillary pericytes within the cortex do not express *α*-SMA, which is thought to be a necessary prerequisite for contraction. It remains unclear which of these vessel types has the greatest impact on CBF regulation, and this formed the basis for our investigations.

Of particular interest given our fluid mechanics approach, were reports of decreases in blood velocity following vasoconstriction^5,19^ or increases in velocity following vasodilation^14,21,45^. According to Poiseuille’s law, which relates blood pressure and the volumetric flow rate to vessel diameter, a decrease in vessel diameter should induce a localised velocity increase in order to conserve mass, rather than the decreases observed *in vivo*. Such a response is expected even when allowing for variation in localised blood flow. These paradoxical observations suggested to us that the observed vasomodulations could be related to changes occurring on the scale of the network, and which could be meaningfully investigated with a large-scale numerical modelling approach.

Alongside vessel diameter changes, we were also interested in probing alternative mechanisms that have been proposed for regulating CBF. For example, Wei et al. ^45^ suggest that erythrocytes act as O_2_ sensors and can alter their shape in order to modify blood flow and vascular O_2_. Questions remain regarding how erythrocytes can actively deform, if erythrocyte modulation couples with arteriolar hyperaemia and if cognitive decline in various diseases is a result of impaired PO_2_-mediated capillary hyperaemia. However, combining each of these mechanisms in a comprehensive mathematical model allowed us to study them in much greater detail than could be achieved in a conventional experimental setting.

Our mathematical modelling provided additional, detailed insights into volumetric flow, haematocrit and PO_2_ (oxygen partial pressure) changes within blood vessels and cortical tissue. These have enabled us to form a clearer understanding of the mechanisms underpinning CBF regulation, which we present here, and which have implications for the interpretation of future *in vivo* experiments.

## Results

### Simulation of blood flow and oxygen delivery in the mouse cortex

Our study used *in vivo* imaging data from the mouse cortex, acquired with a two-photon microscope (Praire Technologies). We acquired image data from adult male SMA-mCherry transgenic mice^2^, which enabled cortical *α*-SMA on the surface of arterioles to be detected via the fluorescence of mCherry (800 nm excitation). Cascade Blue Dextran (800 nm excitation) was also injected intravenously, to allow the simultaneous imaging of blood vessel lumen. Image volumes had dimensions of 420 × 420 × 400 *μm*^3^ and resolution of 2.4 × 2.4 × 4 *μm*^3^ (see Fig 1 a, b). During post-processing, we segmented blood vessel lumen and smooth muscle regions using in-house software, and converted the data into a graph format consisting of nodes and connecting segments. These data provided the geometrical inputs for our mathematical models. Each vessel had an associated diameter and a flag denoting the presence or absence of smooth muscle coverage (see Fig 1 c).

**Figure 1:**
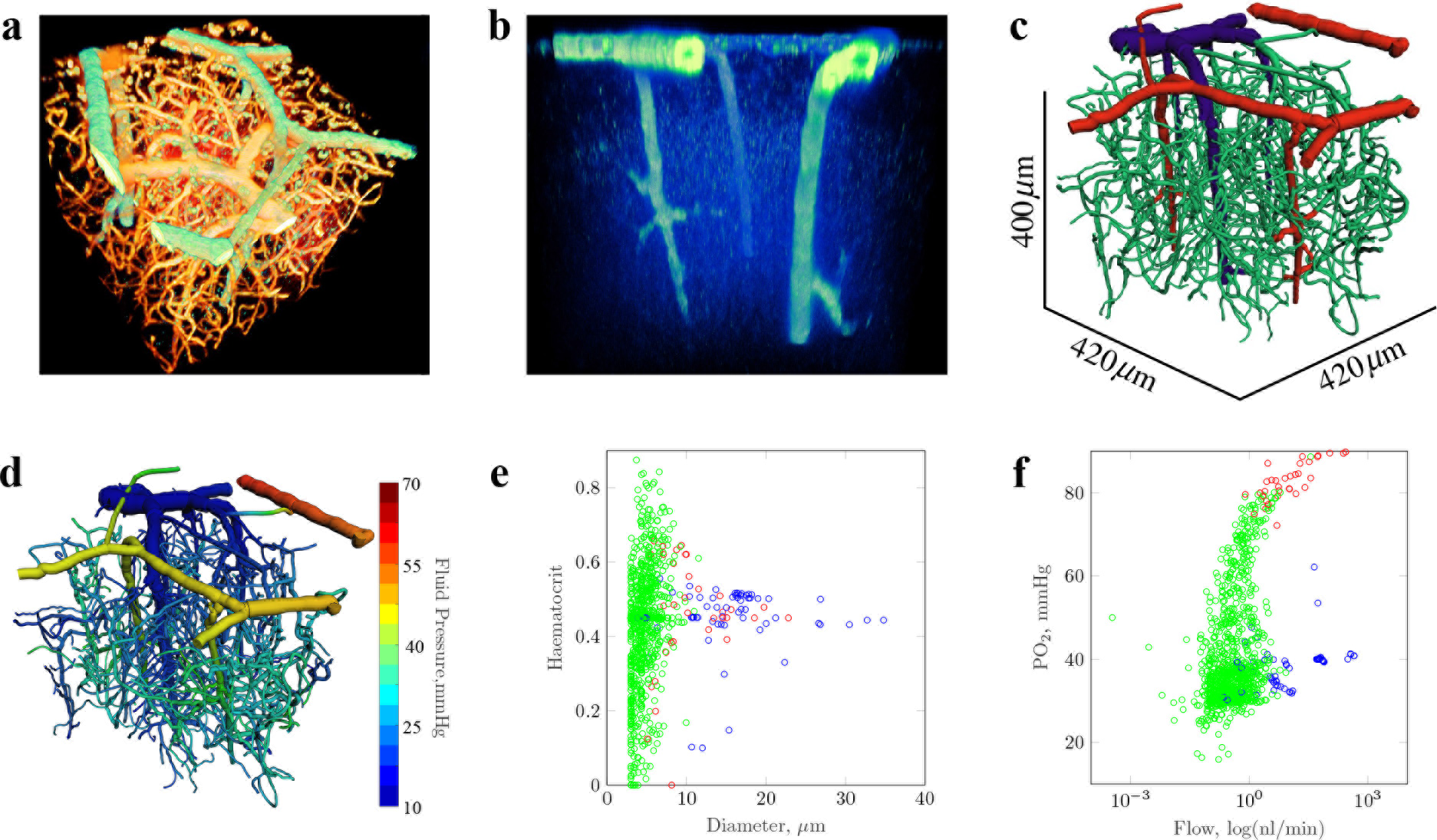
Mathematical modelling of blood flow in the cortical microvasculature, using 3D confocal imaging data from transgenic mice. (a) A view of the cortical vascular network showing smooth muscle actincovered vessels (SMA, green), from transgenic expression of m-Cherry, and blood vessel lumen (orange), from exogeneous administration of flouorescent dextran. (b) An enhanced view showing the location of SMA expressing vessels. (c) The vascular network, following image segmentation, with vessels colour-coded based on classification (red: SMA-expressing arterioles; blue: veins; green: capillaries). (d) Simulated cortical blood pressures within the cortical network. (e) A scatter plot of vessel diameter against simulated, blood vessel haematocrit from our baseline simulation. (f) A scatter plot of volumetric blood flow (nL/min) against intravascular PO_2_ (mmHg). Data points in (e) and (f) are colour-coded according to vessel classification, as per (c).

Our mathematical model was written in C++, in which Poiseuille flow was modelled through individual vessel segments, and was coupled to a Green’s function model for intravascular and tissue oxygen transport ^44^. Our model enabled a complex, quantitative assessment of blood flow within large cortical networks, and included the non-Newtonian effects of blood and haematocrit heterogeneity through application of empirical laws^29^. Boundary conditions for the model were assigned according to previously published computational data^23,24^. Diameter-dependent pressure conditions were assigned to all pial boundaries, with inlet haematocrit set to 0.45, allowing flow solutions to be calculated that minimised the deviation from target pressures and vessel wall shear stresses ^6^. Similarly, boundary PO_2_ was assigned based on experimental data^9,36^, thereby enabling physiologically-accurate solutions.

Our first simulation run provided a “baseline” flow and O_2_ solution, using vascular network architecture derived from imaging data (see Fig 1 d, e, f). This returned a vascular perfusion of 147.1 ml/min/100g, which is in good agreement with previous studies ^26,46^, whilst mean tissue PO_2_ (partial oxygen pressure) was calculated as 29.3 mmHg (with a computed oxygen extraction fraction of 0.47). Flow velocities, mean PO_2_ and SO_2_ (oxygen saturation) were also in the range of previous experimental measurements ^36,45^ and computational studies^12^ (see Table 1). This agreement between our simulation results and literature values gave us confidence that our model could provide physiologically-realistic solutions.

**Table 1:**
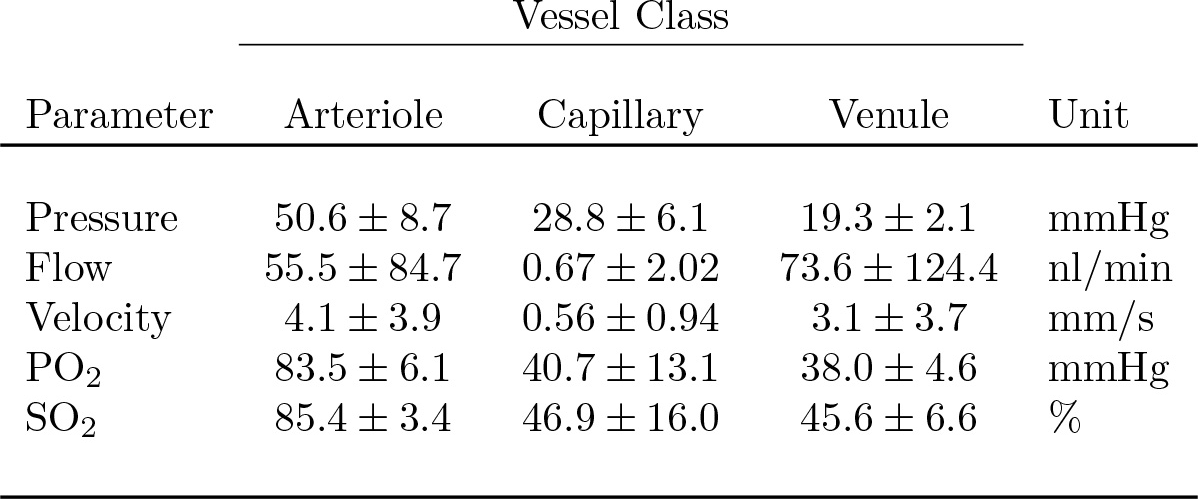
Blood flow and oxygenation summary statistics, for the baseline simulation (data are mean ± standard deviation).

### Optogenetically-induced constrictions in arteriolar smooth muscle propagate to upstream arterioles and induce blood velocity decreases

Hill et al. ^19^ performed single-cell mural constrictions using confocal laser line-scanning to activate ChR_2_ (channelrhodopsin 2). Single-cell constrictions, in which individual mural cells are activated, induced a reduction in vessel diameter. In order to prevent proximal vessel constrictions via the confocal laser, targeted two-photon stimulation of ChR_2_ within small areas of interest along single vessels, was also used to measure the resultant blood velocity modulations at the site of constriction. As a second validation of our model, we performed an *in silico* experiment to replicate this type of intervention.

Following Hill et al. ^19^, constrictions were simulated in three vessel types: 1) penetrating arterioles (smooth muscle coverage and a diameter >10 *μm*), 2) precapillary arterioles (smooth muscle coverage and diameter < 10 *μm*, at the arteriolar/capillary interface) and 3) capillaries (no smooth muscle coverage, and first branching order from the arteriolar/capillary interface). Consistent with those measured by Hill et al. ^19^, we reduced vessel diameter by −19.41, −11.79 or −0.1% (representing peak diameter changes for vessel types 1-3, respectively), in a series of 25 simulations each containing a localised, single-cell constriction (10 *μm* in length).

For single-cell constriction of penetrating and precapillary arterioles, we found a large increase in blood velocity of 53.0 ± 1.6 and 27.5 ± 2.8%, respectively, within the constricted vessel. This increase in velocity at the point of constriction is to be expected, according to fluid dynamical models, and is analogous to constricting a hose pipe, in which fluid velocity increases to conserve mass. However, this conflicts with the observations by Hill et al. ^19^, and others ^5^, who observed a velocity decrease following vessel constriction. This presents a paradox that we sought to resolve.

During functional hyperaemia, vessel dilations can be propagated upstream from the initial site of activation^3^. We hypothesised that optogenetic SMC activation could result in a cascade of vasoconstriction in its local vicinity. We therefore explored three scenarios in addition to single-cell constriction (scenario 1 - see Fig 2); constriction of a single-cell initiating a cascade of constriction: 2) extending to the nearest vessel branching points (single vessel constriction); 3) upstream to the nearest penetrating arteriole and then propagating in both directions perpendicular to the cortical surface (bi-directional cascade); and 4) upstream to the nearest penetrating arteriole and then propagating upstream towards the cortical surface (uni-directional cascade). For consistency, these cases were also investigated for capillary constrictions.

**Figure 2:**
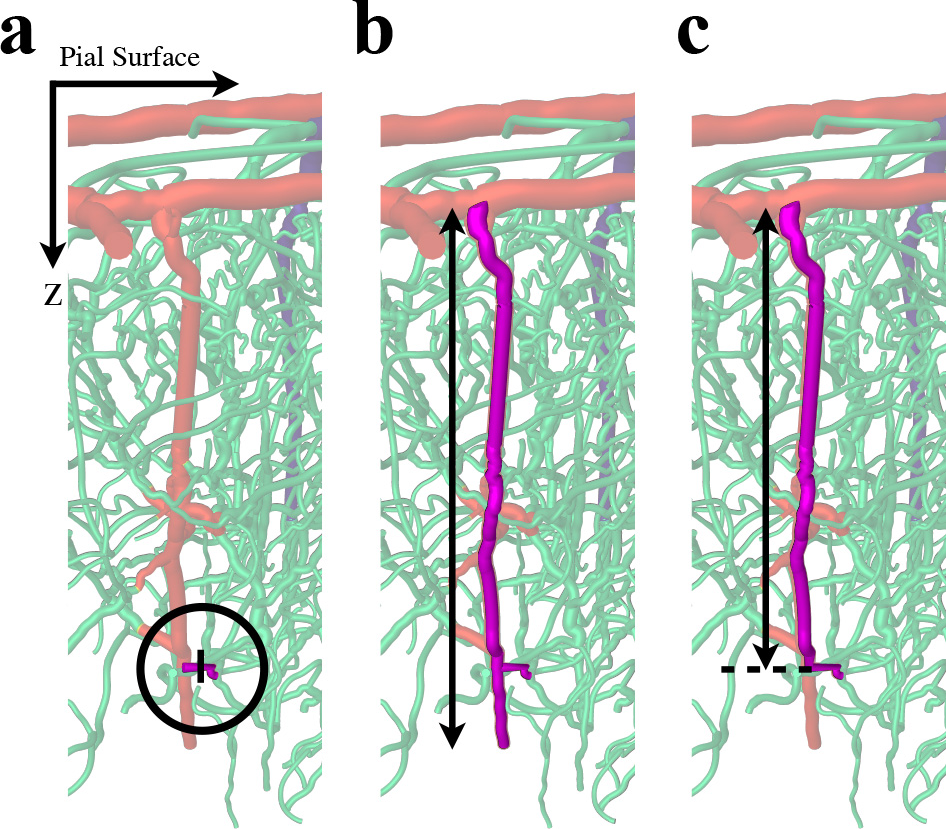
Sites of simulated blood vessel constrictions in SMA-expressing vessels (purple). (a) The location of single branching-order precapillary arteriole constriction (circled); the black line indicates the site of a single-cell constriction. (b) Constriction of a precapillary arteriole, with bi-directional constriction cascade to the entire adjoining penetrating arteriole. (c) Constriction of a precapillary arteriole and uni-directional propagation to upstream sections of the penetrating arteriole. The dashed line indicates the point at which the penetrating arteriole bifurcates to the precapillary arteriole.

We found that, for bi-directional propagation, mean blood velocity increased in penetrating arterioles (see Fig 3 a). In the case of uni-directional propagation, the constricted vessel exhibited a velocity increase of 25.9 ± 12.4%, but unconstricted, downstream vessels exhibited a velocity decrease of 11.7 ± 8.0%, which is consistent with *in vivo* experiments ^5,19^.

**Figure 3:**
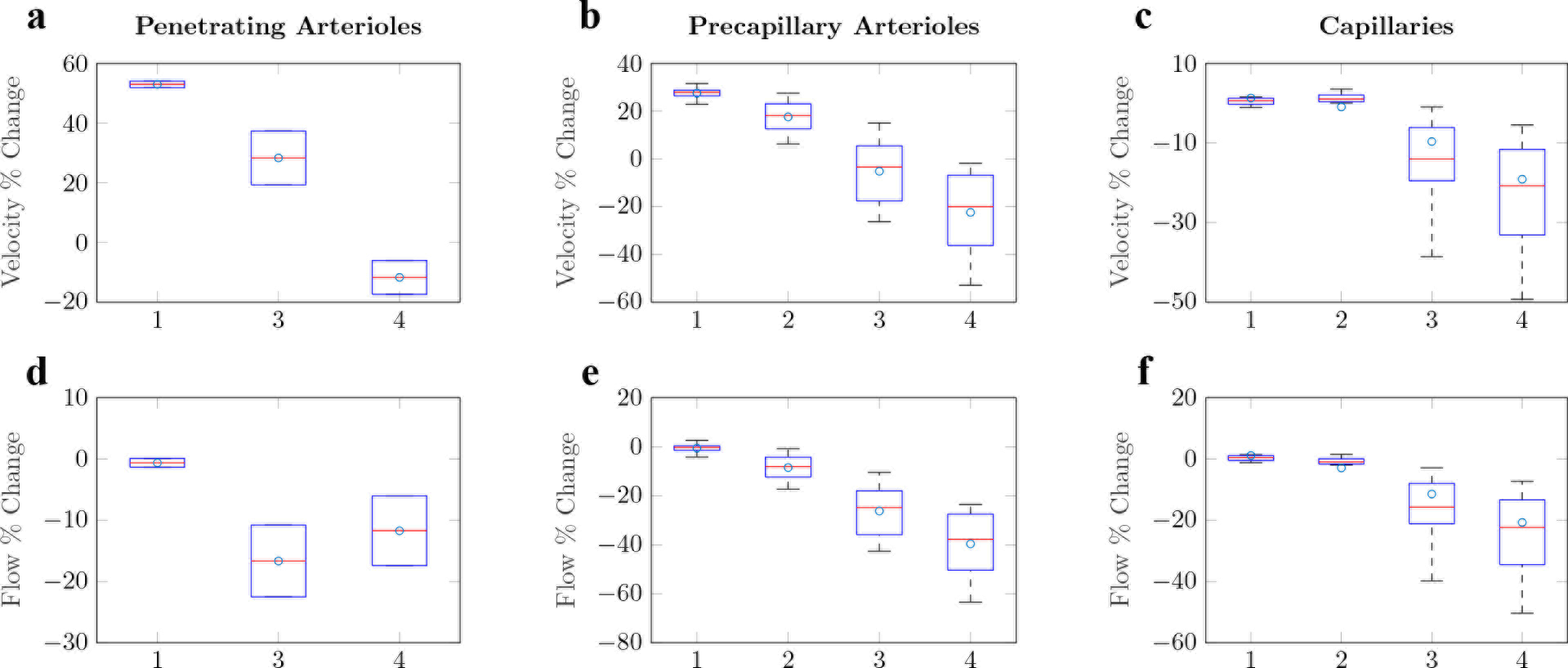
Box plots showing simulated blood velocity and flow changes in response to vasoconstriction. The top row of graphs shows blood velocity changes as a percentage of the baseline solution, and the bottom row shows blood flow changes. From left to right, plots show the responses to constrictions in penetrating arterioles (left), precapillary arterioles (middle) and capillaries (right). Results are presented for each constriction scenario 1) single-cell constriction, 2) constriction of the entire vessel, 3) bi-directional and 4) uni-directional constriction of the local penetrating arteriole. Note, cases 2) and 3) are equivalent for penetrating arterioles and the mean values for each are indicated with circles. Case 4 for penetrating arterioles provides the velocity change for the neighbouring downstream vessel from the site of constriction.

We observed blood velocity changes of 17.6±6.6% and −5.2±14.4% in constricted precapillary arterioles for scenarios 1 (single vessel constriction) and 2 (bi-directional cascade), respectively (see Fig 3 b). Similarly, uni-directional propagation caused blood velocity in constricted precapillary arterioles to decrease by 22.2±8.0%, a magnitude of change that is more consistent with that found *in vivo* ^19^, and so we discounted scenarios 1 and 2. This outcome suggests that optogenetically-induced constrictions of SMA-covered precapillary arterioles propagates upstream along its local penetrating arteriole. These physiologically-plausible results resolve the paradoxical *in vivo* observations of a decrease in blood velocity following blood vessel constriction. It is only by significantly increasing the resistance to blood flow by propagating diameter reductions to a much larger section of upstream vessels, that we observe a velocity decrease in response to vessel constriction. It is also plausible that this mechanism can explain similarly paradoxical increases in blood flow observed following induced vessel dilations of cerebral arterioles.

### Optogenetically-induced constriction of capillary pericytes does not induce constriction propagation

Our simulations found that constriction of capillary pericytes produced a blood velocity increase of 1.3±3.4% within the constricted vessel. This is consistent with *in vivo* observations that capillary diameter changes contribute minimally to changes in blood velocity^19^. For consistency, uni- and bi-directional constriction cascades were also investigated for capillary constrictions.

When constrictions were initiated across single capillary branches, we found blood velocity decreased by 1.0 ± 8.3%. As with single-cell constriction, minimal changes in blood velocity occurred (see Fig 3 c). As pericytes are known to span along entire vessels ^16,19^, whole vessel constriction is arguably a more physiologically representative intervention than the 10 *μm* ‘single-cell’ activation. Inducing bi- and unidirectional constriction cascade, passing from perictyes to SMA-covered vessels, produced velocity decreases far in excess of the variations observed *in vivo* ^19^ (−9.6 ± 21.1 and −18.6 ± 25.3%, respectively - see Fig 3 c). As such, our results suggest that optogenetically-induced constriction of capillaries, via capillaries, does not induce a cascade of constriction along local SMA-covered arterioles.

### The relationship between vasoconstriction and haemodynamic changes is nontrivial

In our simulations, following single vessel constriction, blood flow in penetrating and precapillary arterioles decreased by 0.6 ± 1.0 and 0.6 ± 2.2%, which further decreased to −39.5 ± 13.2% when uni-directional constriction cascade was introduced (see Fig 3 d, e). However, blood velocity showed changes in the opposite direction, with an increase of 53.0 ± 1.6 and 17.6 ± 6.6% for penetrating and precapillary arterioles, and a change of 35.9 ± 12.4 (−11.7 ± 8.0 downstream of constriction) and 22.2 ± 16.9% when uni-directional propagation was introduced. Mathematically, these opposite changes in blood flow and blood velocity observed within penetrating arterioles are a distinct possibility (see Section S1). Therefore, we advise caution against the use of blood velocity as a proxy measurement for flow.

Within capillaries, this non-trivial relationship between blood velocity and flow becomes more complex due to the increased influence of flow resistance instigated by the restricted passage of erythrocytes (see Fig S13). Our simulations of constrictions in SMA-covered vessels, initiated by pericyte constriction, showed that reductions in blood flow and velocity were mediated by decreases in blood viscosity and blood pressure, compared to pressure alone in SMA-covered vessels (see Tables S6 and S7). Since we have shown that optogenetic-induced constriction of pericytes does not induce a cascade of constriction along vessels, these data suggest that dynamically altering blood viscosity within capillaries may provide the most optimal means to regulate capillary hyperaemia.

### Nano-scale, single-capillary pericyte constrictions have a variable and spatially heterogeneous effect on tissue PO_2_

Having investigated the effect of vasoconstriction on blood flow and velocity, we turned our attention to the impact of these changes on intravascular and tissue PO_2_.

The recent study by Kisler et al. ^21^ demonstrated that pericyte loss-of-function weakens capillary ability to modify CBF following neuronal stimuli, leading to neurovascular uncoupling and reduced oxygen supply. It was suggested that the inhibited ability of a capillary to dilate following pericyte dysfunction relates to tissue PO_2_ regulation. However, Hill et al. ^19^ observed minimal vasomotion in pericyte-covered capillaries (a 0.08 ± 0.18% change in diameter) following optogenetic-activation and, in addition, we have shown that at peak constriction, only small changes were exhibited in both blood velocity and flow (unless constriction cascade is introduced). Nonetheless, it is unclear whether these small changes in flow are sufficient to impact cortical oxygenation.

Using our flow constriction solutions, we therefore simulated changes in O_2_ transport within the cortical network. We observed changes in capillary PO_2_ of −0.6 ± 0.08, −7.8 ± 5.9 and −9.1 ± 6.7% for constrictions in penetrating arterioles, precapillary arterioles and capillaries, respectively (which included constriction cascade in SMA-covered vessels - see Fig 4 a, b, c). Tissue PO_2_ changed by −13.0 ± 2.4, −2.7 ± 0.8% and 0.45 ± 0.31%, respectively (see Fig 5 a, b, c).

**Figure 4:**
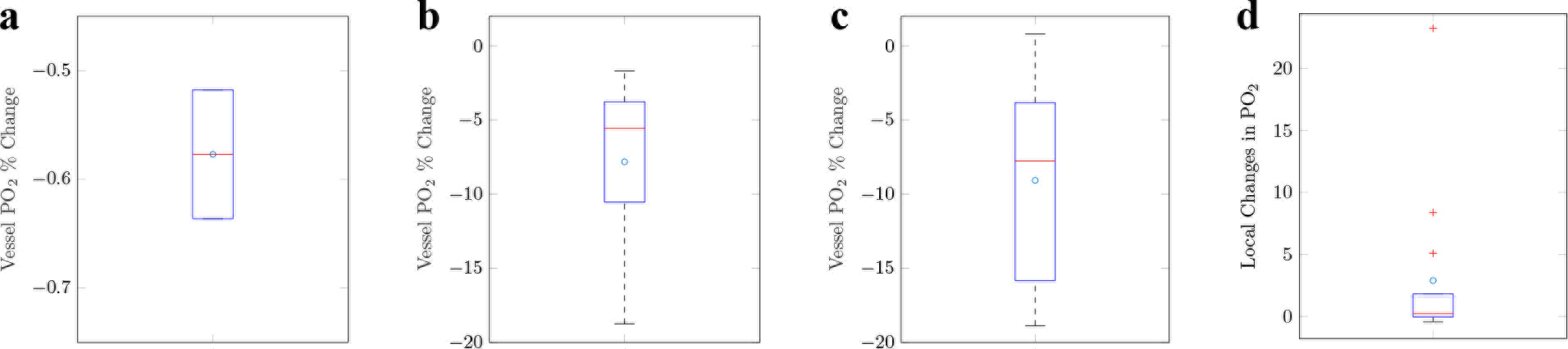
Box plots of simulated vessel PO_2_ changes, compared to baseline, in response to vessel constriction in (a) penetrating arterioles, (b) precapillary arterioles, (c) capillaries, and (d) tissue PO_2_ changes local to constricted capillaries. Outliers (data larger than *q*_3_ + 3(*q*_3_ − *q*_1_)/2 or smaller than *q*_1_ − 3(*q*_3_ − *q*_1_)/2, where *q*_1_ and *q*_3_ are the 25^*th*^ and 75^*th*^ percentiles, respectively) are indicated by red crosses.

**Figure 5:**
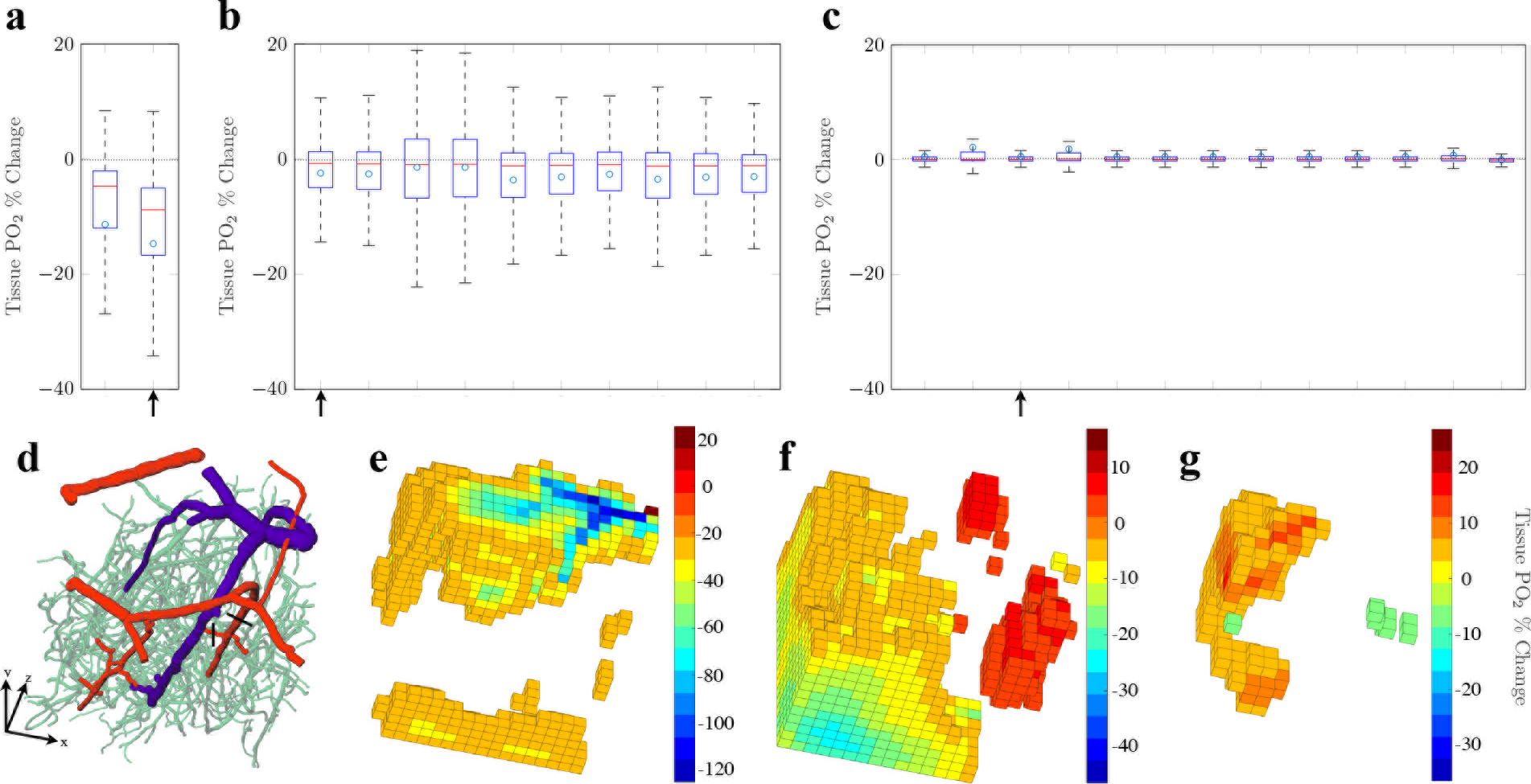
Simulated tissue PO_2_ changes, compared to baseline solutions, in response to blood vessel constriction (a total of 25 constrictions). Box plots are shown of the percentage change in cortical tissue PO_2_ for constrictions in (a) penetrating arterioles, (b) precapillary arterioles and (c) capillaries. Arrows indicate simulation results displayed in (e, f, g). (d) 3-dimensional visualisation of the cortical network, in which black lines indicate the initial site of constriction for (e) and (f), and with the capillary constriction for (g) occurring in the downstream vessel, one branching order from (f). 3-dimensional visualisations of discretised tissue PO_2_ percentage changes, for constrictions in (e) a penetrating arteriole, (f) a precapillary arteriole and (g) a capillary. Note, tissue was discretised into 8000 cubic regions and PO_2_ % changes, with absolute values < 20% for (e) and absolute values < 5% for (f) and (g) are not shown, for clarity.

This suggests that, even with minimal flow changes, significant reductions in PO_2_ were found following nano-scale capillary vasoconstriction, and were similar in size to those exhibited following constrictions in precapillary arterioles. This effect is possible due to the combined effect of small flow reductions alongside vessel haematocrit reductions (−2.9±8.2 and −2.36±6.58%, respectively), the latter of which plays a much larger role in capillaries than in larger vessels and, in the case of micron-scale dilations, has been hypothesised to be an efficient mechanism that can locally alter the distributions of RBCs in microvascular networks^40^.

Nano-scale constriction of capillaries produced a highly variable PO_2_ response, with a heterogeneous distribution of both increases and decreases in PO_2_ in the vicinity of a constriction. For example, in a 60 × 60 × 60 mm^3^ tissue volume centred on the constricted capillary, tissue PO_2_ increased by 2.9 ± 6.6% (rather than an expected decrease; see Fig 4 d), due to the redistribution of haematocrit to neighbouring capillaries.

In addition, we found a strong correlation between changes in blood flow and capillary PO_2_, which was not evident between changes in blood velocity and capillary PO_2_ (see Fig S15, and Table S3). This suggests that blood flow is a more significant factor in determining PO_2_ than blood velocity. The *in silico* study by Liicker et al. ^25^ also found that haematocrit had a larger influence on tissue PO_2_ than RBC velocity in lower order capillaries. In combination, this identifies the limitations in drawing inferences about cortical oxygenation from measurements of blood velocity.

### Arteriolar constrictions induces stable intravascular capillary and tissue oxygenation changes

We have shown that nano-scale constriction of capillaries leads to a significant reduction in vessel PO_2_, yet causes an unpredictable alteration to tissue PO_2_ in the local vicinity, due to haematocrit redistribution. In addition, optogenetic activation of capillary pericytes does not appear to induce constriction cascade to local SMA-covered arterioles. However, due to the complex molecular mechanisms surrounding pericyte activation^14,15^, under normal physiological conditions, this effect could occur via other signalling pathways, which may then induce more stable and predictable regulation of local tissue PO_2_ surrounding capillaries. Based on this hypothesis, we investigated the effects of simultaneous constriction of a capillaries with unidirectional constriction cascade to local SMA-covered vessels. The results of these simulations were then compared with those in which capillaries remain unconstricted, but constrictions were propagated to SMA-covered vessels.

Our results show that local constriction of SMA-covered vessels had a similar, if not greater, impact on capillary haemodynamics than when coupled to capillary constriction (see Fig 6 a, b, c, d, e). SMA constriction caused intravascular capillary PO_2_ at unconstricted sites to reduce by 9.1±7.3%, led by a greater reduction in capillary flow and haematocrit (30.0 ± 11.5 and 29.2 ± 30.1%, respectively). In comparison, constriction cascade to SMA-covered vessels, coupled with capillary constriction, reduced PO_2_, flow and haematocrit by a similar or diminished amount: 9.1 ± 6.7, 20.7 ± 24.9 and 25.5 ± 26.8%, respectively.

**Figure 6:**
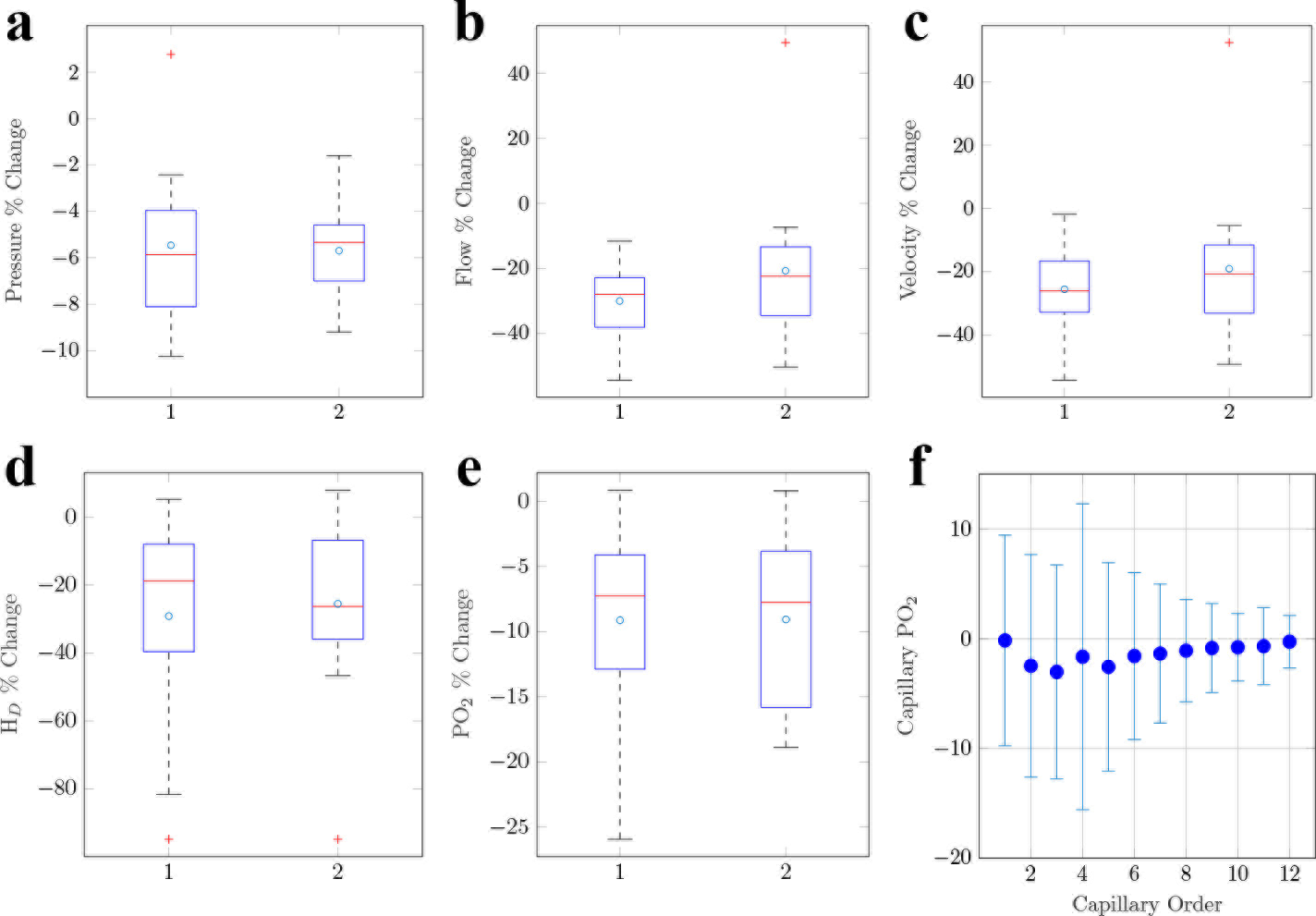
Simulating SMA constriction cascade following pericyte constriction. Percentage changes in (1) unconstricted and (2) constricted capillaries with uni-directional SMA propagation for (a) pressure, (b) flow, (c) velocity, (d) haematocrit (H_*D*_) and (e) vessel PO_2_. (f) A plot of mean change in capillary PO_2_ against capillary order, as a result of local SMA constriction. Error bars represent standard deviation and, consequently, the redistribution of O2 through the cortical network.

We next analysed the impact of SMA constriction cascade on each capillary branching order, which showed that local SMA cascade in the absence of pericyte-led constriction can effectively alter downstream capillary intravascular PO_2_ (see Fig 6 e, f) in addition to reducing tissue PO_2_. These results suggest that SMA constriction cascade, led by precapillary arterioles, does not necessarily need to be coupled to pericyte capillary constriction in order to actively regulate the passage of intravascular and interstitial O_2_. Thus, our simulations suggest that SMA-covered vessels have the capacity, in isolation, to regulate blood flow and oxygenation in a targeted and predictable manner throughout the cortex.

### Functional erythrocyte deformability has the capacity to drive capillary hyperaemia

We have demonstrated that constriction via smooth muscle dynamically alters blood flow, velocity, fluid pressure and PO_2_. Precapillary arterioles and capillaries adjacent to penetrating arterioles offer the greatest contribution to hydraulic resistance^12^, which implicates alternative mechanisms for regulating capillary hyperaemia other than through changes in blood vessel diameter. One such mechanism was proposed by Wei et al. ^45^, who suggested that erythrocytes can deform in response to low vessel PO_2_, causing a reduction in blood viscosity^45^ (but without initially altering blood haematocrit).

This formed the basis for our next set of simulations, in which our baseline cortical flow solution identified capillary vessels that exhibited PO_2_ lower than 25 mmHg (1.7% of all capillary segments), these became sites for spontaneous erythrocyte deformation (instantaneous elongation). We reduced blood viscosity in these vessels by 20% to mimic the effects of erythrocyte deformation.

We found that spontaneous erythrocyte deformations increased blood velocity, on average, by 9.8 ± 6.2% (with equivalent changes in flow). This is consistent with the peak velocity changes measured empirically by Wei et al. ^45^, and was accompanied by an average vessel PO_2_ increase of 12.9 ± 24.3%. This provides support for the hypothesis that spontaneous erythrocyte deformation is a plausible mechanism for increasing vessel PO_2_. Given that we have shown that SMA-covered arterioles can dynamically alter capillary PO_2_, we performed a further simulation to investigate the combined effect of arteriolar diameter changes, alongside erythrocyte deformation. We induced a diameter dilation of 4.5%^45^ to all SMA-exhibiting vessels (excluding pial vessels) alongside erythrocyte deformation in capillaries with low-oxygen tolerance (as defined above). On average, velocity and flow increased in these capillaries by 10.7±5.7%, and PO_2_ increased by 18.2±31.3%. This result shows that global arteriolar dilation coupled to erythrocyte deformation amplified the PO_2_ change in these capillaries on average by 41.1%, compared to isolated deformation.

We also found that a large subset (97.4%) of capillaries exhibited a marked increase in blood flow and PO_2_ in response to erythrocyte deformation, both when modelled in isolation, and when erythrocyte deformation was coupled to arteriolar dilation (see Fig 6 b and 7). This could limit the efficacy of erythrocytes for providing targeted, isolated changes in tissue oxygenation that are not compromised by neighbouring regions. However, whether such effects have a significant impact on neurovascular coupling would require further investigation. Likewise, whilst our simulations strongly implicate SMA-covered vessels as the main modulatory apparatus for cortical oxygenation, they are unable to determine the mechanism that triggers such a response from the capillary level. Rather, our simulations indicate that both erythrocyte deformation and pericyte signalling are both plausible mechanisms, and worthy of experimental investigation.

**Figure 7:**
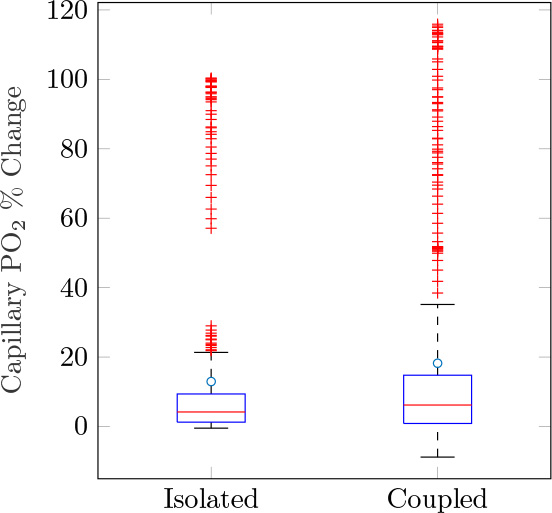
Capillary PO_2_ changes initiated by erythrocyte deformation. Box plots show the percentage change in PO_2_ for capillaries initially below 25 mmHg, for erythrocyte deformation occuring in isolation, and when coupled to arteriolar (SMA) dilation.

## Discussion

The regulation of cortical blood flow has been studied extensively, yet details of the mechanisms underpinning its precise control remain elusive. This is due in part to the multiplicity of potential influences, which are interdependent and challenging to isolate empirically. We therefore sought to develop an original paradigm, composed of both *in vivo* imaging and mathematical modelling, to explore cortical microcirculatory blood flow and oxygen transport regulation mechanisms in a block of mouse cortex, imaged using a two-photon microscope *in vivo*.

The mathematical models which constitute our *in silico* platform have been thoroughly tested and validated in previous computational studies in order to accurately simulate blood flow^6,7,8,9,13,23,24,33^ and oxygenation^7,43,44^ in physiological microvascular networks. In addition, our platform has been parametrised and the results compared against a wealth of experimental data^4,5,9,19,26,28,36,37,45,46^ in order to validate its simulations.

A key advantage of our modelling approach is its ability to provide complete network flow, haematocrit and oxygen distributions, simultaneously, which would be impossible to obtain in a conventional experimental context. This also allowed us to perform novel, controlled *in silico* experiments on a real-world cortical network structure. By recreating experimental designs from the literature, *in silico*, such as by Hill et al. ^19^ and Wei et al. ^45^, we have demonstrated the ability of our paradigm to reproduce physiologically realistic solutions and provided further physiological insights. In contrast to previous studies, our study combined *in vivo* genetic cell labelling and mathematical modelling to probe cortical microvascular structure and function, which are otherwise unavailable in isolation.

Hill et al. ^19^ provided an excellent start point, as this study provided detailed *in vivo* measurements of blood velocity in multiple vessel types, following vessel constriction. They demonstrated that activation of SMC-covered arterioles in mouse cortex caused a significant decrease in diameter (−19.41 ± 1.9%) and blood velocity (−44.62 ± 11.8%), whereas blood vessels covered by pericytes showed no significant change in either diameter or velocity (−0.08 ± 0.18% and −2.22 ± 3.16%, respectively). This reported decrease in blood velocity in response to constriction of SMA-covered arterioles (using optogenetic stimulation) presented a paradox, given that Poiseille’s law predicts that, by decreasing vessel diameter, blood velocity should increase in order to conserve mass transport, even with marginal changes in inflow (in contrast, with an increase in vessel diameter, blood velocity should decrease). Our simulations explain this paradox if this localised activation induces a cascade of upstream constrictions, from precapillary arterioles to descending penetrating arterioles. The increased flow resistance induces a velocity decrease of similar magnitude to that measured experimentally in Hill et al. ^19^. This propagation mechanism has previously been demonstrated by Chen et al. ^3^ in the instance of arteriolar dilation, and our simulations provide strong support for this effect in the case of arteriolar constriction.

Invoking constriction propagation enabled us to resolve the paradoxical observations of blood velocity decrease in response to vasoconstriction, but also demonstrated a non-trivial relationship between changes in blood velocity (directional speed) and blood flow (volume transfer per unit time) following vasoconstriction. Conversely, flow was strongly correlated with capillary oxygenation. Poiseuille’s law forms the basis of our model and is often invoked in the literature to link changes in blood velocity to blood flow through a single vessel. However, we have shown that this is not always the case as it does not incorporate the network flow response, nor the redistribution of haematocrit. We have shown that it is possible for a vessel to exhibit an increase in blood velocity, post-constriction, whilst observing a reduction in flow. Therefore, caution is needed when inferring flow changes from measured blood velocity changes; blood velocity is a poor proxy for flow or oxygenation.

Constriction propagation was not necessary to explain Hill et al. ^19^’s velocity changes in response to capillary pericyte activation. When we simulated constriction cascade from capillaries, velocity decreases were far in excess of *in vivo* measurements. However, we cannot infer whether this is a nuance of optogenetic activation, or if constriction cascade cannot be propagated from capillary pericytes. Likewise, isolated capillary constrictions induced minimal blood flow changes, compared with arteriolar constrictions.

It is accepted that smooth muscle cells control arteriolar hyperaemia, however the mechanisms controlling capillary hyperaemia are debated^14,19,21^ and the role of pericytes remains controversial. The capacity of pericytes to modulate cerebral blood flow has been measured *in vivo* through vasoconstriction^19^ and vasodilation^14,21^, with data (inferred from velocity measurements) suggesting that capillaries contribute minimally to variations in CBF^19^. Conversely, pericyte dysfunction has also been linked to neurovascular uncoupling^21^. Our simulations showed that nano-scale constrictions are able to significantly alter capillary and tissue PO_2_, although the resultant changes were spatially heterogeneous, with constriction-induced velocity decreases accompanied by both increases and decreases in nearby capillaries. This variation across the network was due to the redistribution of blood flow and haematocrit to neighbouring vessels. Interestingly, flow and oxygenation changes caused by constriction cascade from capillaries to arterioles were similar if not smaller than when propagation of arterioles emanated solely from local precapillary arterioles.

Our results therefore suggest that arterioles have the capacity to regulate capillary hyperaemia without the need for pericyte-led capillary constriction. Hall et al. ^14^ suggested that, due to their proximity to neurons, neuronal signalling to pericytes occurs before signalling to smooth muscle. Additionally, Kisler et al. ^21^ demonstrated that pericyte-deficient mice had a diminished oxygen supply to the cortex. As such, we hypothesise the pericytes may purely act as signalling conduits to upstream arteriolar smooth muscle, allowing for a localised vasomotion response by arterioles. Any pericyte-led neurovascular uncoupling may inhibit signalling to local arterioles and thereby reduce the ability to regulate cortical O_2_. In summary, our results suggest that arterioles have the capacity to dynamically modulate both arteriolar and capillary hyperaemia in a coordinated and stable fashion.

However, other (potentially complimentary) regulation mechanisms have also been proposed in the literature. We investigated the recent hypothesis that erythrocytes act as O_2_ sensors which regulate their own deformability in order to increase flow velocity in response to low PO_2_ ^45^. We reduced blood viscosity in vessels below a PO_2_ threshold in order to mimic the drop in flow resistance observed experimentally^45^. The subsequent velocity changes as a result of erythrocyte deformation correlated well with peak velocity changes seen *in vivo* ^45^ and flow increases resulted in significant PO_2_ increases within selected capillaries with further increases exhibited when coupled to global arteriolar dilation. This provides support for erythrocyte deformation as a plausible mechanism for regulating vascular PO_2_ and cerebral blood flow. However, this functional response requires further empirical investigation to determine, for example, the scale of erythrocyte deformation and its ability to dynamically alter tissue PO_2_ (which can then be more precisely modelled).

The limited extent of the network used in this study meant that the effects of vascular steal from adjoining vascular networks could not be studied in detail, and larger networks could provide further insights. It would be of interest to investigate the influence of erythrocyte deformability incorporated to facilitate O_2_ regulation under pathophysiological conditions^45^. Likewise, capillaries have been observed to dilate in response to micro-occlusions^35^, potentially suggesting a more dynamic response via the vascular network than we were able to simulate here. Each of these effects can be further explored in future studies, placing our modelling platform in an ideal position to perform detailed hypothesis-testing.

We have established a novel platform which combines *in vivo* cortical microvascular data with computational models to gain insights into cerebral blood flow regulation and oxygen delivery. This platform has enabled a thorough understanding of mural cell contribution to cortical haemodynamics, and the formulation of hypotheses that had not arisen from the *in vivo* work in isolation. For example, we have proposed that optogenetic stimulation of SMA-covered arterioles induces a cascade of upstream constrictions and that such activation of pericytes does induce an equal response. Also nano-scale constrictions of pericyte covered capillaries can significantly alter capillary PO_2_, although these changes are spatially heterogenous. In comparison, arterioles have the capacity to hyperaemia without the need for pericyte-led capillary constriction, suggesting pericytes may purely act as signalling conduits to upstream arteriolar smooth muscle. Moving forwards, we anticipate that the findings from this work will give rise to new experimental and mathematical modelling research to further our understanding of neurovascular coupling in both health and disease.

## Methods

### Animal Experiments

All animal procedures were approved by the Institutional Animal Care and Use Committee (IACUC) at Yale University, with all methods performed in accordance with the relevant guidelines and regulations. We acquired our images from adult male SMA-mCherry transgenic mice^2^ aged 120 days housed in a 12:12 light:dark cycle animal vivarium with food and water provided ad libitum. We performed acute craniotomies to acquire *in vivo* two-photon fluorescence based microvascular three-dimensional reconstructions. Briefly animals were anaesthetised via intraperitoneal injections of ketamine (100 mg/kg) and xylazine (10 mg/kg) anaesthetic and a 3 mm craniotomy was performed over the somatosensory cortex. The underlying dura was removed and a #0 cover glass was placed over the craniotomy. To visualise the microvasculature, Cascade Blue Dextran (10,000 mw, ThermoFisher Cat# D1976) was injected intravenously prior to imaging. Images were acquired on a two-photon microscope (Prairie Technologies) equipped with a mode-locked MaiTai laser (Spectra Physics) with a 20x water immersion objection (Zeiss 1.0 NA). Excitation of Cascade Blue Dextran and mCherry in SMA-mCherry mice was achieved using 800 nm two-photon excitation. A full outline of the methods used is outline in Hill et al. ^19^.

### Network Segmentation

Image segmentation was undertaken through thresholding, where we set threshold levels by visual inspection to isolate high intensity fluorophore signals from background noise, for both dextran and mCherry (SMA) signals. For dextran signals, discontinuities within the network, mostly due to imaging artifacts and absent dextran within vessel lumen, were corrected manually in Amira (FEI, Oregon, USA). Thresholded data were thinned using a skeletonisation algorithm^39^ and converted to a segment and node format in Amira. We determined SMA coverage, for each vessel, by the distance of each blood vessel centerline to mCherry-positive voxels using in-house software written in IDL (Harris Geospatial Solutions, Boulder, Colorado, USA). If positive signal was detected within 1.5 vessel radii, over at least 90% of a vessel’s length, then we categorised that vessel as positive for SMA coverage. This process enabled reconstruction of the cortical microvasculature with vessel classification based on SMA expression, as shown in Fig 1.

A summary of network statistics (for example, vessel diameters, lengths and classifications) in shown in Table S2. We segmented the vascular network into a series of interconnected nodes and segments, where each segment was assumed to be a cylindrical tube. The final network consisted of 26,662 segments, connected by 25,678 nodes, of which 388 were boundary nodes. As venules were not classified by SMA expression, non-SMA vessels above a threshold of 8 *μm* were identified and those forming a connected tree-like structure were categorised as venules.

### Mathematical Blood Flow Model

The mathematical model was coded in C++ and is comprised of the discrete network flow model of Pries et al. ^32^ alongside the flow estimation algorithm of Fry et al. ^6^, where the structural properties of the segmented network and hemodynamic parameters are used as inputs. Here we presented the mathematical models followed by a section on boundary condition (BC) and parameter assignment. The discrete-network blood flow model has been thoroughly tested using mesenteric networks ^28^ in which blood flow measurements were taken in individual vessels ^6,29,30,32^ with equivalent models applied to cortical networks ^13,23,24^.

For our cortical microvascular network, no flow or pressure conditions are known at boundaries of the tissue. The method by Fry et al. ^6^ extends the discrete network flow model of Pries and Secomb ^29^ to networks in which BCs are unknown, based on the concept that the microcirculation is regulated in response to signals relating to flow and shear stresses ^31^. The scheme estimates unknown BCs by minimising the squared deviation from specified target network wall shear stresses and pressures values derived from independent information about typical network hemodynamic properties. In essence, the algorithm removes the need to define conditions at all boundary nodes, to one where simulation sensitivity is weighted towards the definition of these two target parameters.

Our segmented microvascular network (see Fig 1) is represented by a series of vessel segments connected by nodal junctions or, in the case of boundary nodes, one-segment nodes which form a boundary to the microvascular network. We defined a positive flow direction from the start node to end node of each vessel segment. Under the assumption of Poiseuille flow and conserving flow at blood vessel junctions, the relationship between nodal pressures, *p*_*k*_ and the boundary boundary fluxes *Q*_0*i*_ is given by

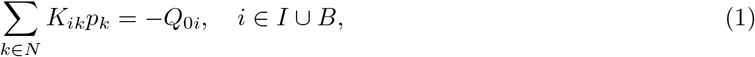

where *N* is the set of all nodes, *I* is the set of all interior nodes and *B* is the set of all boundary nodes with known BCs. For all interior nodes, conservation dictates that *Q*_0*i*_ = 0, however, if *i* is a known boundary node, *Q*_0*i*_ is the inflow (or outflow if negative). Note, at least one pressure BC needs to be assigned for a unique solution. The matrix *K*_*ik*_ represents network conductivity

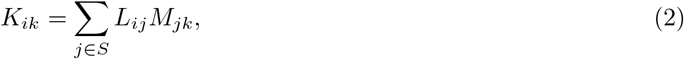

where *S* is the set of all segments,

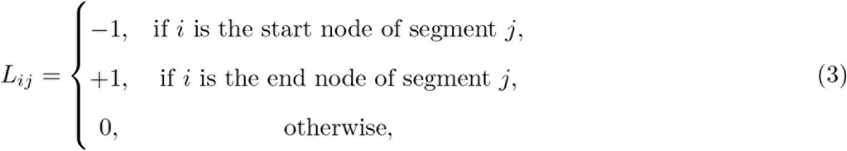

and

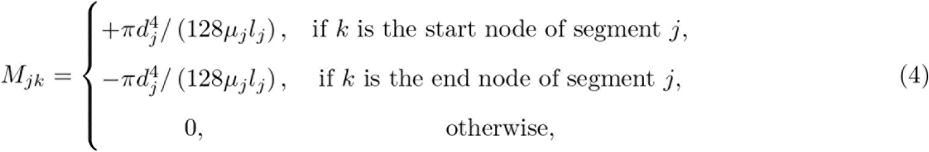

is the matrix of vessel conductances where *l*_*j*_, *d*_*j*_ and *μ*_*j*_ denote the length, diameter and effective viscosity of segment *j*, respectively.

We used empirical *in vivo* blood viscosity laws, which prescribe the effective viscosity as a function of vessel diameter and haematocrit, to compute *μ*_*j*_ and consequently incorporate non-Newtonian effects in each individual microvessel ^29^. Network haematocrit heterogeneity plays an important part in network flow resistance; however, access to data on haematocrit distributions is limited. In the absence of these data, empirical descriptions can be incorporated to include red blood cell screening, the process by which the balance of fluid forces determines a blood cell’s destination^27^. The approach requires an iterative procedure in which flow *Q*_*j*_ in vessel *j* is used to update haematocrit, *H*_*Dj*_ in each vessel. Here, boundary haematocrit was assigned as 0.45 and the process was repeated until the sequence converges to given tolerances (10^−3^) for both flow and haematocrit.

In the absence of measured flow data, the ill-posed system, (1), requires further assumptions to obtain a unique solution. The method proposed by Fry et al. ^6^ sought to solve a constrained optimisation problem, formulated in terms of a Lagrangian objective function that is defined by

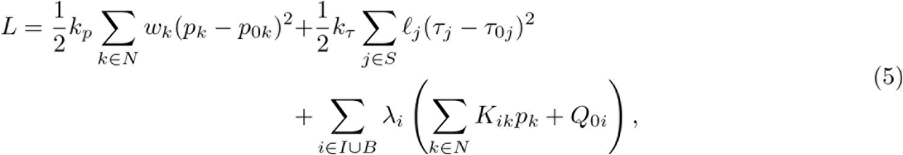

where *p*_0*k*_ is the target pressure at node *k*, *τ*_*j*_ is the wall shear stress in segment *j*, *τ*_0*j*_ is the corresponding target shear stress, *k*_*p*_ and *k*_*τ*_ are weighting factors associated with the pressure and shear deviations from the target values, λ_*i*_ is the Lagrange multiplier associated with node *i* and *w*_*k*_ is the vessel length associated with node *k*. Setting *dL*/*dp_i_* = 0 and combing with equation equations (1) yields the following sparse linear system

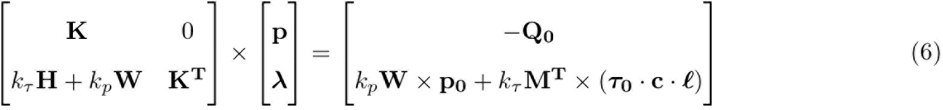

where **K**, **W**, **M** and **H**, denote the matrix forms of *K*_*ik*_, *w*_*k*_, *M*_*ij*_ and *H*_*ik*_, respectively. Note, **W** is a diagonal matrix with entries *w*_*k*_ and *H*_*ik*_ is defined as

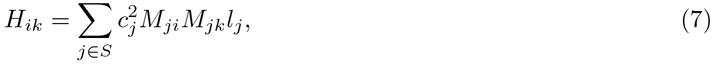

where 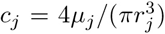. The remaining variables **p**, **λ**, **Q**_0_, **p**_0_, ***τ***_0_, **c** and **ℓ** are the vector forms of *p*_*k*_, λ_*k*_, *Q*_0*i*_, *p*_0*k*_, *τ*_0*j*_, *c*_*j*_ and *ℓ*_*j*_ respectively. The resulting sparse square linear system, (6), contains *n*_*I*_ + *n*_*B*_ + *n*_*N*_ linear equations, where *n*_*I*_, *n*_*B*_ and *n*_*N*_ are the number of interior, known boundary and the total number of nodes. We solved this system for unknowns *p*_*i*_ and *λ*_*i*_ using standard numerical methods.

### Mathematical Oxygen Transport Model

We used our flow solutions from the discrete-network model to parametrise the following well-established approach which describes steady-state intravascular and tissue oxygen transport, whereby tissue is represented as a homogeneous medium with oxygen diffusivity *D* and solubility *α*, as outlined in a series of papers ^10,20,44^.

Derived from Fick’s Law, tissue PO_2_, *P*, conservation satisfies

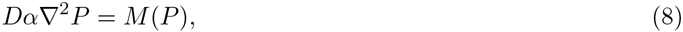

where *M*(*P*) is the oxygen consumption rate. The dependence of oxygen consumption on PO_2_ is represented by a Michaelis-Menten relationship

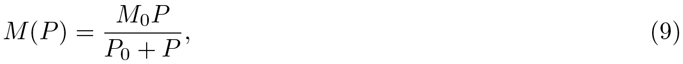

where *M*_0_ is a uniform oxygen demand in the tissue given a non-limiting supply of oxygen and *P*_0_ represents the PO_2_ at half-maximal consumption.

The rate of convective oxygen transport along a vessel segment is given by

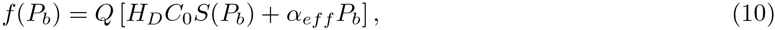

where *P*_*b*_ is the blood PO_2_ level, *Q* is the blood flow rate, *H*_*D*_ is the discharge haematocrit, *C*_0_ is the concentration of haemoglobin-bound oxygen in a fully saturated erythrocyte, *S* is the oxyhemoglobin saturation and *α*_*eff*_ is the effective solubility of oxygen in blood. The Hill equation presents a simple description of oxyhemoglobin saturation, as a consequence of oxygen binding to hemoglobin, and is given by

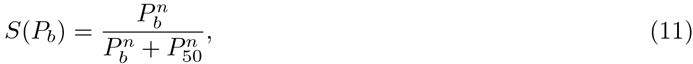

where *n* is the Hill exponent and *P*_50_ is the PO_2_ at 50% saturation. The effective solubility of oxygen in blood is given by

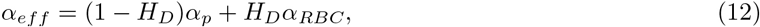

where *α*_*p*_ and *α*_*RBC*_ are the solubilities in blood plasma and in RBCs, respectively. These values are similar^18^ implying haematocrit has a small weighting towards *α*_*eff*_. Thus, allowing us to approximate the effective oxygen solubility by a constant value.

Conservation of oxygen flux infers

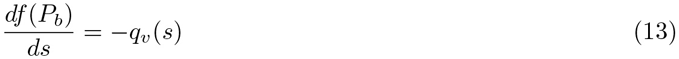

in each vessel segment, where *s* defines segment length and *q*_*v*_ is the rate of diffusive oxygen efflux per unit of *s*.

Continuity of oxygen flux and PO_2_ (under the assumption of cylindrical vessel segments) across the vessel walls yields

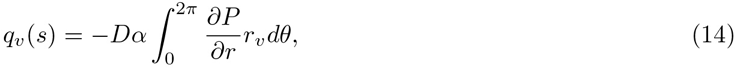

where *r*_*v*_ is the vessel radius and integration is performed over the circumference of the vessel, denoted by the azimuthal angle *θ*. The blood vessel delivering the oxygen generally exceeds local PO_2_ levels at the interface with the surrounding tissue. Hellums ^17^ defined the following jump condition at the interface

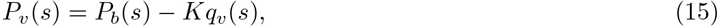

where *P*_*v*_(*s*) is the average tissue *PO*_2_ and *K* represents intravascular resistance to radial oxygen transport.

The oxygen transport model given by (8)-(15) is solved numerically^41^ by formulating it in terms of Green’s functions, following the established approach reported in Secomb et al. ^44^, whereby blood vessels are represented as a set of discrete oxygen sources. Using superposition principles, the resulting fields from these sources represent the PO_2_ field in the tissue. If the rate of oxygen uptake within the tissue is prescribed, the only unknowns in the problem are the strengths of both the sources and sinks. We apply this mathematical model to our cortical microvascular network.

Modelling oxygen transport, the Green’s function, *G*(x; x′), for a given tissue domain may be defined as the PO_2_ at a point x resulting from a unit point source at x′, through

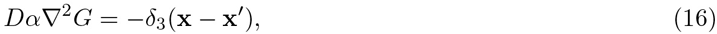

where *D* is oxygen diffusivity, *α* is oxygen solubility and *δ*_3_ is the delta function in three-dimensions. Solving the adjoint problem the *PO*_2_ is thus given by

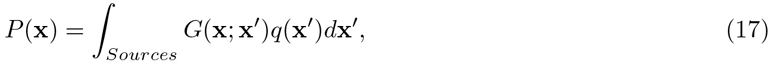

where *q*(x) represents the distribution of source strengths.

The oxygen field resulting from a blood vessel can be represented by a series of sources down its centre. Following the approach of Hsu and Secomb ^20^, the oxygen sources are located along the vessel axis and uniformly distributed around the circumference of the vessel on the blood-tissue boundary. Whilst the approach uses an iterative scheme to incorporate the non-linear reaction rate kinetics, resolving the spatial oxygen fields for complex microvascular networks comes at a relatively low cost when compared to finite-difference methods alongside minimising artefacts associated with assigning BCs ^20,44^. The numerical implementation of the Green’s model is outlined in Secomb et al. ^44^. In addition, a recent extension incorporates time-dependent solute transport ^42^.

## Boundary Conditions & Parameter Assignment

Sensitivity analysis equivalent to that investigated by Fry et al. ^6^ estimation model on the mesenteric networks of Pries et al. ^28^ was replicated. Increasing the number of applied BCs reduced the normalised root mean square deviation between segment flows predicted with partial boundary information and those simulated when all BCs (*Q*_0*j*_) are known. As network boundary pressure and flow data were unavailable for our cortical network, we extrapolated a function to compute arteriolar or venular pressures based on the vessel diameters using data from previous intra-cortical blood flow simulations by Lorthois et al. ^23^,^24^ (see Fig S10 a). Lorthois et al. ^24^ simulated a steep pressure drop on the arteriolar side compared to venules (a drop of 75 to 15 mmHg, with a mean capillary pressure of 31 mmHg), an asymmetry consistent with previous experimental studies ^22,32^. A function was fitted to this data in order to apply pressure conditions to the boundaries of our pial arteriolar and venula vessels (representing 2.3% of all BCs in the network - see Fig S9), using vessel classifications shown in Fig 1 c. In the absence of measured data, we applied a haematocrit of 0.45 at all boundary nodes.

In the network flow estimation scheme, we assumed the target pressures at each node and target shear stresses in each segment to be constants (equal in magnitude for all segments). Target pressure, *p*_0*j*_, assigned as the mean capillary pressure computed by Lorthois et al. ^24^, 31 mmHg. The sign of the shear stress, *τ*_0*j*_, in each segment is equal to the corresponding flow sign, but this cannot be known until we obtain a solution. The target shear stresses (initially set with random flow signs and to a value of 5 dyn/cm^2^) are used to estimate the flow distribution and then updated in each segment for the next iterative step, whilst updating the value of *τ*_0*j*_ to the mean vessel target wall shear stress on the previous iteration. During optimisation (with application to the rat mesentery networks of Pries et al. ^28^), Fry et al. ^6^ initially set *k*_*τ*_ = 10^−4^ and on each iteration the value of *k*_*τ*_ was doubled until two consecutive iterations gave the same flow directions in all segments. The constant *k*_*p*_ is used to maintain pressures in a physiologically-realistic range (tested against the measured flows in the mesenteric networks ^6^) and was arbitrarily set to 0.1 in order to bias the nodal pressures towards the specified target pressure. We employed an identical approach in this study.

When simulating constriction propagation of vessels, we used baseline capillary boundary pressures as BCs. Here, pressure was assumed to be maintained by adjoining tissue blocks. Similarly, the arteriolar and venular baseline flow solutions were used as BCs, with the exception that the base of the local penetrating arteriole was set as an unknown BC. We explored a variety of BC assignment in order to produce the required physiological velocity changes, in addition to network response. For example, inspired by observations of venular distension in response to arteriolar dilation^4^, we investigated venular compression to match blood velocity changes to constriction, compared to those *in vivo*.

Simultaneous BCs for oxygen transport cannot be determined from available experimental information. In the Green’s function method, the tissue volume is considered as being embedded in an infinite domain with the same diffusivity, where no oxygen sources or sinks are located externally to the given tissue region. Hence, we formed a well-posed problem without imposing explicit BCs on the outer surface of the tissue region. The method itself is parametrised by the discrete-network model, by inputting the computed flow and haematocrit values for each vessel segment. Alongside this, the model requires us to assign boundary PO_2_ for network inlets, which can be a combination of arterioles, capillaries and venules. Based on experimental and simulated data, we assigned arterioles and venules PO_2_ values of 90 and 40 mmHg, respectively^9^. In order to assign heterogenous capillary PO_2_, we formed a data-fitted curve by extrapolating experimental data^36^ in which PO_2_ was measured relative to cortical depth (see Fig S14). The model’s intravascular and tissue oxygen transport parameter values are given in Table S8. Compared to previous cortical oxygenation studies^44^, we assigned the consumption parameter, *M*_0_, based on a recent study of CMRO_2_ estimation (mean value of 1.71 *μ*mol cm^−3^ min^−1^ used^37^). In this study, the Krogh cylinder model was applied, whilst taking into account that the rate of oxygen consumption is approximately 80-85% of its maximum value^1,11^.

## Acknowledgements

We would like to thank R.A. Hill and J. Gruztendler for acquiring and supplying the imaging data. In addition, Seyyed Haqshenas and Iva Burova for discussions into the oxygen transport model parameter assignment.

### Author Contributions

All authors were involved in the design of the study. S.W.S processed and segmented the image data. P.W.S performed numerical simulations. All authors contributed to the writing of the manuscript.

### Competing Interests

We have no competing interests.

## References

[1] Angleys, H., Østergaard, L. and Jespersen, S. N. 2015, ‘The effects of capillary transit time heterogeneity (CTH) on brain oxygenation’, Journal of cerebral blood flow and metabolism: official journal of the International Society of Cerebral Blood Flow and Metabolism 35(5), 806–17.

[2] Armstrong, J. J., Larina, I. V., Dickinson, M. E., Zimmer, W. E. and Hirschi, K. K. 2010, ‘Characterization of bacterial artificial chromosome transgenic mice expressing mCherry fluorescent protein substituted for the murine smooth muscle α-actin gene’, Genesis 48(7), 457–463.

[3] Chen, B. R., Kozberg, M. G., Bouchard, M. B., Shaik, M. A. and Hillman, E. M. C. 2014, ‘A Critical Role for the Vascular Endothelium in Functional Neurovascular Coupling in the Brain’, Journal of the American Heart Association 3(3).

[4] Drew, P. J., Shih, A. Y. and Kleinfeld, D. 2011, ‘Fluctuating and sensory-induced vasodynamics in rodent cortex extend arteriole capacity.’, Proceedings of the National Academy of Sciences of the United States of America 108(20), 8473–8.

[5] Fernández-Klett, F., Offenhauser, N., Dirnagl, U., Priller, J. and Lindauer, U. 2010, ‘Pericytes in capillaries are contractile in vivo, but arterioles mediate functional hyperemia in the mouse brain’, Proceedings of the National Academy of Sciences of the United States of America 107(51), 22290–22295.

[6] Fry, B. C., Lee, J., Smith, N. P. and Secomb, T. W. 2012, ‘Estimation of Blood Flow Rates in Large Microvascular Networks’, Microcirculation 19, 530–538.

[7] Fry, B. C., Roy, T. K. and Secomb, T. W. 2013, ‘Capillary recruitment in a theoretical model for blood flow regulation in heterogeneous microvessel networks’, Physiological Reports 1(3).

[8] Gagnon, L., Sakadžic, S., Lesage, F., Mandeville, E. T., Fang, Q., Yaseen, M. a. and Boas, D. a. 2015, ‘Multimodal reconstruction of microvascular-flow distributions using combined two-photon microscopy and Doppler optical coherence tomography’, Neurophotonics 2(1), 015008.

[9] Gagnon, L., Sakadžić, S., Lesage, F., Musacchia, J. J., Lefebvre, J., Fang, Q., Yücel, M. A., Evans, K. C., Mandeville, E. T., Cohen-Adad, J., Polimeni, J. R., Yaseen, M. A., Lo, E. H., Greve, D. N., Buxton, R. B., Dale, A. M., Devor, A. and Boas, D. A. 2015, ‘Quantifying the microvascular origin of BOLD-fMRI from first principles with two-photon microscopy and an oxygen-sensitive nanoprobe’, The Journal of neuroscience: the official journal of the Society for Neuroscience 35(8), 3663–75.

[10] Gagnon, L., Smith, A. F., Boas, D. A., Devor, A., Secomb, T. W. and Sakadžić, S. 2016, ‘Modeling of Cerebral Oxygen Transport Based on In vivo Microscopic Imaging of Microvascular Network Structure, Blood Flow, and Oxygenation’, Frontiers in Computational Neuroscience 10, 82.

[11] Gjedde, A., Johannsen, P., Cold, G. E. and Ostergaard, L. 2005, ‘Cerebral metabolic response to low blood flow: possible role of cytochrome oxidase inhibition’, Journal of cerebral blood flow and metabolism: official journal of the International Society of Cerebral Blood Flow and Metabolism 25(9), 1183–96.

[12] Gould, I. G., Tsai, P., Kleinfeld, D. and Linninger, A. 2017, ‘The capillary bed offers the largest hemodynamic resistance to the cortical blood supply’, Journal of cerebral blood flow and metabolism: official journal of the International Society of Cerebral Blood Flow and Metabolism 37(1), 52–68.

[13] Guibert, R., Fonta, C. and Plouraboué, F. 2010, ‘Cerebral blood flow modeling in primate cortex’, .Journal of cerebral blood flow and metabolism: official journal of the International Society of Cerebral Blood Flow and Metabolism 30(11), 1860–1873.

[14] Hall, C. N., Reynell, C., Gesslein, B., Hamilton, N. B., Mishra, A., Sutherland, B. a., O’Farrell, F. M., Buchan, A. M., Lauritzen, M. and Attwell, D. 2014, ‘Capillary pericytes regulate cerebral blood flow in health and disease’, Nature 508(7494), 55–60.

[15] Hamilton, N. B., Attwell, D. and Hall, C. N. 2010, ‘Pericyte-mediated regulation of capillary diameter: a component of neurovascular coupling in health and disease’, Frontiers in neuroenergetics 2(May), 1–14.

[16] Hartmann, D. A., Underly, R. G., Grant, R. I., Watson, A. N., Lindner, V. and Shih, A. Y. 2015, ‘Pericyte structure and distribution in the cerebral cortex revealed by high-resolution imaging of transgenic mice.’, Neurophotonics 2(4), 041402.

[17] Hellums, J. D. 1977, ‘The resistance to oxygen transport in the capillaries relative to that in the surrounding tissue’, Microvascular research 13(1), 131–6.

[18] Hellums, J. D., Nair, P. K., Huang, N. S. and Ohshima, N. 1995, ‘Simulation of intraluminal gas transport processes in the microcirculation’, Annals of Biomedical Engineering 24(S1), 1–24.

[19] Hill, R. A., Tong, L., Yuan, P., Murikinati, S., Gupta, S. and Grutzendler, J. 2015, ‘Regional Blood Flow in the Normal and Ischemic Brain Is Controlled by Arteriolar Smooth Muscle Cell Contractility and Not by Capillary Pericytes’, Neuron 87(1), 95–110.

[20] Hsu, R. and Secomb, T. W. 1989, ‘A Green’s function method for analysis of oxygen delivery to tissue by microvascular networks’, Mathematical Biosciences 96(1), 61–78.

[21] Kisler, K., Nelson, A. R., Rege, S. V., Ramanathan, A., Wang, Y., Ahuja, A., Lazic, D., Tsai, P. S., Zhao, Z., Zhou, Y., Boas, D. A., Sakadžić, S. and Zlokovic, B. V. 2017, ‘Pericyte degeneration leads to neurovascular uncoupling and limits oxygen supply to brain’, Nature Neuroscience (20), 406–416.

[22] Lipowsky, H. H. n.d., ‘Microvascular rheology and hemodynamics’, Microcirculation (New York, N.Y.: 1994) 12(1), 5–15.

[23] Lorthois, S., Cassot, F. and Lauwers, F. 2011a, ‘Simulation study of brain blood flow regulation by intra-cortical arterioles in an anatomically accurate large human vascular network: Part I: Methodology and baseline flow’, NeuroImage 54(2), 1031–1042.

[24] Lorthois, S., Cassot, F. and Lauwers, F. 2011 b, ‘Simulation study of brain blood flow regulation by intra-cortical arterioles in an anatomically accurate large human vascular network. Part II: Flow variations induced by global or localized modifications of arteriolar diameters’, NeuroImage 54(4), 2840–2853.

[25] Lücker, A., Secomb, T. W., Weber, B. and Jenny, P. 2017, ‘The relative influence of hematocrit and red blood cell velocity on oxygen transport from capillaries to tissue’, Microcirculation 24(3), e12337.

[26] Maeda, K., Mies, G., Oláh, L. and Hossmann, K. A. 2000, ‘Quantitative measurement of local cerebral blood flow in the anesthetized mouse using intraperitoneal [14C]iodoantipyrine injection and final arterial heart blood sampling’, Journal of cerebral blood flow and metabolism: official journal of the International Society of Cerebral Blood Flow and Metabolism 20(1), 10–4.

[27] Pries, A. R., Ley, K., Claassen, M. and Gaehtgens, P. 1989, ‘Red cell distribution at microvascular bifurcations’, Microvascular research 38(1), 81–101.

[28] Pries, A. R., Ley, K. and Gaehtgens, P. 1986, ‘Generalization of the Fahraeus principle for microvessel networks’, The American journal of physiology 251(6 Pt 2), H1324–32.

[29] Pries, A. R. and Secomb, T. W. 2005, ‘Microvascular blood viscosity in vivo and the endothelial surface layer’, American journal of physiology. Heart and circulatory physiology 289(6), H2657–H2664.

[30] Pries, A. R. and Secomb, T. W. 2011, Blood Flow in Microvascular Networks, in ‘Microcirculation’, 2nd edn, John Wiley & Sons, Inc., Hoboken, NJ, USA, pp. 3–36.

[31] Pries, A. R., Secomb, T. W. and Gaehtgens, P. 1995, ‘Structure and hemodynamics of microvascular networks: heterogeneity and correlations’, The American journal of physiology 269(5 Pt 2), H1713–22.

[32] Pries, A. R., Secomb, T. W., Gaehtgens, P. and Gross, J. F. 1990, ‘Blood flow in microvascular networks. Experiments and simulation’, Circulation Research 67(4), 826–834.

[33] Pries, A. R., Secomb, T. W., Gessner, T., Sperandio, M. B., Gross, J. F. and Gaehtgens, P. 1994, ‘Resistance to blood flow in microvessels in vivo.’, Circulation research 75(5), 904–915.

[34] Reichold, J., Stampanoni, M., Lena Keller, A., Buck, A., Jenny, P. and Weber, B. 2009, ‘Vascular graph model to simulate the cerebral blood flow in realistic vascular networks.’, Journal of cerebral blood flow and metabolism: official journal of the International Society of Cerebral Blood Flow and Metabolism 29(8), 1429–1443.

[35] Ren, H., Du, C., Yuan, Z., Park, K., Volkow, N. D. and Pan, Y. 2012, ‘Cocaine-induced cortical microischemia in the rodent brain: clinical implications’, Molecular Psychiatry 17(10), 1017–1025.

[36] Sakadžić, S., Mandeville, E. T., Gagnon, L., Musacchia, J. J., Yaseen, M. a., Yucel, M. a., Lefebvre, J., Lesage, F., Dale, A. M., Eikermann-Haerter, K., Ayata, C., Srinivasan, V. J., Lo, E. H., Devor, A. and Boas, D. a. 2014, ‘Large arteriolar component of oxygen delivery implies a safe margin of oxygen supply to cerebral tissue’, Nature Communications 5, 5734.

[37] Sakadžic, S., Yaseen, M. A., Jaswal, R., Roussakis, E., Dale, A. M., Buxton, R. B., Vinogradov, S. A., Boas, D. A. and Devor, A. 2016, ‘Two-photon microscopy measurement of cerebral metabolic rate of oxygen using periarteriolar oxygen concentration gradients’, Neurophotonics 3(4), 045005.

[38] Sanderson, C. and Curtin, R. 2016, ‘Armadillo: a template-based C++ library for linear algebra’, Journal of Open Source Software Vol. 1, p.26.

[39] Schindelin, J., Arganda-Carreras, I., Frise, E., Kaynig, V., Longair, M., Pietzsch, T., Preibisch, S., Rueden, C., Saalfeld, S., Schmid, B., Tinevez, J.-Y., White, D. J., Hartenstein, V., Eliceiri, K., Tomancak, P. and Cardona, A. 2012, ‘Fiji: an open-source platform for biological-image analysis’, Nature Methods 9 (7), 676–682.

[40] Schmid, F., Reichold, J., Weber, B. and Jenny, P. 2015, ‘The impact of capillary dilation on the distribution of red blood cells in artificial networks’, American Journal of Physiology - Heart and Circulatory Physiology 308(7), H733–H742.

[41] Secomb, T. W. 2011, ‘Green’s function method for simulation of oxygen transport to tissue’. URL: http://physiology.arizona.edu/people/secomb/greens

[42] Secomb, T. W. 2015, ‘A Green’s function method for simulation of time-dependent solute transport and reaction in realistic microvascular geometries’, Mathematical medicine and biology: a journal of the IMA p. 031.

[43] Secomb, T. W., Hsu, R., Beamer, N. B. and Coull, B. M. 2000, ‘Theoretical simulation of oxygen transport to brain by networks of microvessels: effects of oxygen supply and demand on tissue hypoxia’, Microcirculation (New York, N.Y.: 1994) 7, 237–247.

[44] Secomb, T. W., Hsu, R., Park, E. Y. H. and Dewhirst, M. W. 2004, ‘Green’s Function Methods for Analysis of Oxygen Delivery to Tissue by Microvascular Networks’, Annals of biomedical engineering 32(11), 1519–1529.

[45] Wei, H. S., Kang, H., Rasheed, I.-Y. D., Zhou, S., Lou, N., Gershteyn, A., McConnell, E. D., Wang, Y., Richardson, K. E., Palmer, A. F., Xu, C., Wan, J. and Nedergaard, M. 2016, ‘Erythrocytes Are Oxygen-Sensing Regulators of the Cerebral Microcirculation’, Neuron 91(4), 851–862.

[46] Xu, K., Radhakrishnan, K., Serhal, A., Allen, F., LaManna, J. C. and Puchowicz, M. A. 2011, Regional Brain Blood Flow in Mouse: Quantitative Measurement Using a Single-Pass Radio-Tracer Method and a Mathematical Algorithm, Springer US, pp. 255–260.

